# Diverse thermosensory receptors and neurons mediate the neural coding of oral cooling in the mouse trigeminothalamic tract

**DOI:** 10.1101/2022.06.24.497556

**Authors:** Jinrong Li, Christian H. Lemon

## Abstract

Different sets of peripheral and medullary trigeminal neurons respond across a cooling gradient applied to intraoral skin. Here we applied electrophysiology to anesthetized mice to study if different types of cool-driven trigeminothalamic neurons convey oral cooling information to the thalamus. We monitored spiking responses to oral stimulation with cold (≤13°C), cool (21°C to 28°C), neutral (35°C), and warm/hot (≥40°C) water in single trigeminal nucleus caudalis (Vc) neurons physiologically tested for projections to the thalamus. We also recorded oral thermal responses from Vc neurons in mice gene deficient for the cooling and menthol receptor TRPM8 to study afferent mechanisms of central oral thermosensory activity. We found that thalamic-projecting Vc neurons that respond to oral cooling comprise heterogeneous cell types. These cell types showed unique temporal response kinetics across cool and cold temperatures, with tuning to select ranges of a cooling gradient. The combined thermal activity of multiple, differently tuned types of trigeminothalamic cooling neurons offered greater contrast between cold, cool, and warm temperatures in multivariate analysis than the responses of the individual neural types alone, agreeing with a neural population code for cooling information. Compared to control, TRPM8 deficient mice demonstrated a loss of Vc neurons tuned to mild oral cooling, but maintained Vc cells responsive to intense cold. Notably, distinctions between Vc population responses to mild cool and warm temperatures were impaired in TRPM8 deficient mice, suggesting a role for TRPM8 in oral warmth recognition. Diverse receptors and neurons mediate oral cooling signals carried by the trigeminothalamic pathway.

## INTRODUCTION

Oral cooling sensations frequently accompany flavor perception and pleasure experienced during ingestion due to the high resting temperature of intraoral skin (Green, 1986). Cooling and cold sensations inside the mouth also contribute to elements of oral nociception and pain, with, for example, cold-related pain accompanying tooth decay and inflammation (Bernal et al., 2021).

In mammals, cooling stimulation of the oral mucosa excites trigeminal (V) ganglion neurons, which supply somatosensation and nociception to the orofacial region. Several lines of evidence show that in the V ganglion, there are multiple types of primary thermosensory neurons that respond to cooling, defined in part by their sensitivity to select ranges of cooling temperatures. *In vitro* functional data show cool-driven mouse V ganglion cells can be classified as low- or high-threshold based on their sensitivity to a small or comparably larger temperature drop, respectively, from near physiological warm (Bautista et al., 2007; Madrid et al., 2009). Relatedly, functional imaging of mouse V ganglion neurons *in vivo* revealed distinct types of neurons responsive to different ranges of cooling temperatures applied to the oral cavity, including cells predominantly responsive to mild cooling and neurons excited by intense cold (Yarmolinsky et al., 2016; Leijon et al., 2019). Thus, in the periphery, different subsets of cool-driven V afferent fibers can activate across a cooling gradient applied to trigeminal-supplied fields, including intraoral skin.

The fibers of V ganglion neurons partly form and follow the descending trigeminal tract in the brain stem (Capra and Dessem, 1992) to terminate in the trigeminal nucleus caudalis (Vc). Oral cooling excites Vc neurons (Carstens et al., 1998; Zanotto et al., 2007), with our prior neurophysiological data showing cooling-driven mouse Vc cells comprise subgroups responsive to select ranges of mild cool or intense cold oral temperatures (Lemon et al., 2016). This implies that in central V circuits, subpopulations of neurons sensitive to distinct segments of cooling temperatures underly the neural representation of cool and cold oral thermal sensations. This segmentation follows the subtypes of cool-driven V ganglion neurons reported in peripheral studies (Poulos and Lende, 1970b; Yarmolinsky et al., 2016; Leijon et al., 2019). However, whether cool- and cold-active Vc neurons are local circuit interneurons or projection neurons with axons targeting a common downstream brain region was not examined, nor were the orosensory receptor mechanisms that contributed to their thermal responses.

Here we used neurophysiology to study how sensory information about diverse cool and cold oral temperatures is represented by mouse Vc thermosensory neurons with physiologically defined projections to the thalamus. Such Vc cells potentially contribute orofacial thermal information to a thalamocortical pathway. We also studied how genetic silencing of the transient receptor potential (TRP) ion channel TRP melastatin 8 (TRPM8), a cold and menthol receptor expressed by trigeminal fibers (McKemy et al., 2002; Bautista et al., 2007), impacted Vc neural activity to oral temperatures.

We found that cooling sensitive Vc neurons that project to the thalamus include heterogeneous cell types that display diverse tuning and temporal response kinetics to cool and cold stimulation of the oral cavity. The combined activity of these cell types was found to provide neural information that distinguished a broad range of cool and cold temperatures, which agrees with an ensemble, or combinatorial, neural code for cooling (Lemon et al., 2016; Ran et al., 2016; Wang et al., 2018; Leijon et al., 2019; Lemon, 2021). Moreover, silencing TRPM8 resulted in a loss of Vc cells responsive to mild oral cooling but not intense noxious-like cold, implying multiple receptors contribute to neural information about oral cooling represented by V circuits within the brain.

## MATERIALS AND METHODS

### Mice

Forty adult male and female C57BL6/J mice (body weight 20.1 - 32.1 g; stock no. 000664, The Jackson Laboratory) and 21 male and female TRPM8 gene deficient (knockout) mice (body weight 18.6 - 35.6 g; stock no.008198, The Jackson Laboratory) were used. Before recording, all mice were housed in a vivarium that maintained an ambient temperature of ∼23°C and a 12:12-hour light-dark cycle. Food and water were available ad libitum. Animals were prepared for electrophysiological recording using procedures approved by the Institutional Animal Care and Use Committee of the University of Oklahoma and in accordance with the established guidelines set by the National Institutes of Health Guide for Care and Use of Laboratory Animals.

### Surgical preparation

Animals were initially anesthetized before surgery with an intraperitoneal (i.p.) injection of ketamine mixed with xylazine (ketamine:10 mg/ml, xylazine: 1 mg/ml; dosage 10 ml/kg). To reduce bronchial secretions, atropine (24 µg/kg, i.p.) was also administered. Once anesthetized, mice were tracheotomized and a tracheal tube inserted to facilitate breathing during liquid stimulation of the mouth and maintenance of isoflurane gas anesthesia. Mice were secured in a stereotaxic instrument with ear bars (model 930; David Kopf Instruments, Tujunga, CA). The distal end of the trachea tube was positioned inside the tip of custom concentric pressure/vacuum tubing. This tubing allowed anesthetized mice to freely inhale vaporizer-controlled isoflurane (in 100% oxygen gas) and scavenged waste respiratory gas, as described (Li et al., 2020). Anesthesia was maintained during experiments by a continuous flow of 1.2 - 1.5% isoflurane, breathed by mice through the custom system.

Under anesthesia, a rostrocaudal mid-line incision was made on the scalp. Lambda and bregma were exposed and leveled to the same dorsoventral plane using measurements taken with a stereotaxic electrode manipulator arm (model 1460-61, David Kopf Instruments). A small, lateral portion of the occipital bone was removed to expose the dorsal surface of the medulla and allow recording electrode access to the dorsal Vc. A craniotomy was made on the frontal bone above the caudal medial region of the contralateral ventral posteromedial nucleus of the thalamus (VPM) to position a stimulating electrode, which was used to electrophysiologically test Vc neurons for thalamic projections, as below.

A small silk thread was passed behind the lower incisors and lightly drawn taut to deflect the lower jaw downward. Therefore, the tongue could be extended from the mouth by a small rostroventral suture combined with the weight of a small bulldog clamp. The lower incisors were trimmed with rongeurs to not damage the protruding tongue. Heart rate and blood oxygen level were monitored by a pulse oximeter (MouseSTAT^®^ Jr., Kent Scientific, Torrington, CT). Body temperature was maintained at36.5 - 37°C with a heating pad and thermal probe.

### Electrophysiology

As graphically summarized in Figure 1A, we electrophysiologically recorded extracellular action potentials (APs) evoked by oral thermal and chemesthetic stimulation in single Vc neurons in mice. Moreover, we also applied weak electrical current pulses to the oral sensory area of the contralateral VPM via the stimulating electrode to attempt to antidromically backfire the recorded Vc cells from the thalamus. This circuit analysis aimed to identify Vc neurons that maintained thalamic projections and composed the trigeminothalamic tract, presumably indicative of Vc neurons with common function in contributing to thalamocortical processing.

**Figure 1.**
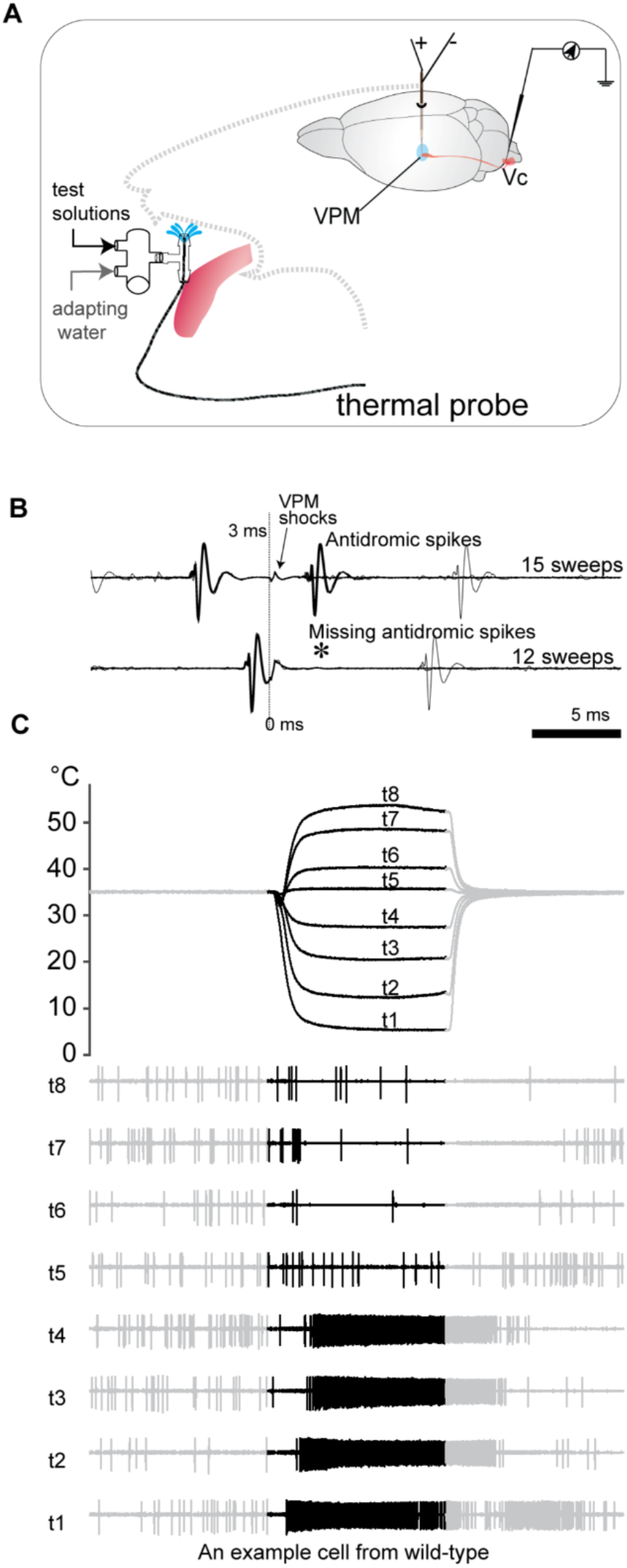
Electrophysiological recording of Vc thermosensory neurons that project to the thalamus in mice. (A) Antidromic and sensory response recording schematic. An antidromic stimulation electrode is unilaterally placed in the orosensory thalamus (VPM). A recording electrode then monitored electrophysiological activity from individual oral thermosensing neurons in the contralateral Vc. (B) Collision test for thalamic projection in a single Vc neuron. Antidromic responses driven by VPM stimulation were triggered at 3 (top) and 0 (bottom) msec following random orthodromic spikes generated by this cell. Antidromic spikes are present (top) but absent (bottom,*) during long- and short-interval stimulation, respectively, indicative of successful spike collision. Thus, this Vc neuron projects to the VPM. (C) Traces track change in oral temperature on 8 oral thermal stimulation trials (top, t1 - 18) and time-locked electrophysiological responses by the Vc neuron in panel B (bottom). This neuron, which projected to the thalamus, responded to cool and cold temperature stimulation of the oral cavity.

The tip of a concentric bipolar stimulating electrode (200 µm O.D.; Microprobes, Gaithersburg, MD) was unilaterally positioned in the VPM using stereotaxic coordinates and electrophysiological guidance. To do this, we first identified general coordinates for the orosensory thalamus by targeting a smaller-diameter tungsten recording microelectrode to the VPM [0.5 - 1.9 mm caudal of bregma, 1.0 - 1.2 mm lateral of the midline, and 3.5 – 4.4 mm ventral to the brain surface (Franklin and Paxinos, 2008)].

The microelectrode was ventrally advanced by an electronic micropositioner (model 2660; David Kopf Instruments) while monitoring for neurons responsive to oral stimuli, including room temperature (cool) water applied to the mouth, as below, or touching/brushing the tongue with a cotton swab. Neural responses to these stimuli confirmed the location of the orosensory VPM. The final VPM coordinates where neurons were found to respond to oral stimuli were then used to target the tip of the concentric bipolar stimulating electrode to the orosensory VPM. After withdrawing the tungsten recording electrode, the bipolar stimulating electrode was ventrally advanced to 200 µm dorsal of the target VPM area. The bipolar electrode was placed into the brain only once to mitigate potential tissue damage.

Once the VPM stimulating electrode was in place, the tip of a tungsten recording microelectrode (3 – 10 MΩ; FHC, Bowdoin, ME) was lowered into the contralateral dorsolateral Vc. The electrode was ventrally advanced using a hydraulic micropositioner (Model MHW-4, Narishige, Tokyo, Japan) attached to a stereotaxic micromanipulator (Model SM-11, Narishige). The probe was advanced in ∼2 µm steps while monitoring for Vc activity to oral cooling stimulation, achieved using oral flow of cool water as below. Neural signals were AC amplified (Grass P511, with high-impedance probe), band-pass filtered (0.3 - 10 kHz), and monitored on an oscilloscope and loudspeaker. Isolated Vc neurons were identified based on waveform consistency. APs were digitally sampled at 25 kHz (1401 interface and Spike2 software; CED, Cambridge, UK) and time-stamped to the nearest 0.1 msec.

Thermosensitive Vc neurons were tested for projections to the orosensory VPM by applying cathodal electrical pulses (100 µA, 0.2 msec) to the VPM bipolar electrode. While doing so, the bipolar electrode was gradually advanced ventrally by ∼100 µm. If VPM pulses could trigger APs in the Vc neuron, the VPM stimulation electrode was moved in 10 µm steps in its tract while identifying the threshold for electrical current stimulation (i.e., the smallest electrical current amplitude applied to the VPM that could trigger APs in Vc cells; thresholds ranged from 60 to 70 µA). Threshold current was used for antidromic circuit testing. As shown by the example in Figure 1B, Vc neurons were considered to show antidromic excitation from the VPM, indicative that VPM pulses excited their axons arriving at the thalamic area, if they displayed 1) a stable evoked response latency, 2) an ability to follow high-frequency (≥ 200 Hz) electrical pulses delivered to the VPM, and 3) collision of orthodromic spikes with putative antidromic spikes (Lipski, 1981). Vc neurons that met all 3 criteria were considered successfully antidromically invaded and to project to the contralateral VPM (Vc^Thal^ neurons). Vc neurons that did not meet all 3 criteria for antidromic invasion were considered to not project to the VPM (Vc^noThal^ neurons).

### Oral stimulation

Delivery of thermal fluids to the oral cavity was accomplished using a custom apparatus, as described for mice (Wilson and Lemon, 2013; Li and Lemon, 2015). In brief, this system allowed for timed delivery to the mouth of temperature-controlled oral stimulus and rinse solutions, and rapid switching between them. Solution flow rates were ∼1.4 mL/sec. Fluid delivered is this manner broadly stimulated oral epithelia, including rostral and caudal regions of the tongue and oral cavity (Wilson and Lemon, 2013).

We recorded AP discharge rates and patterns in single Vc neurons during oral presentation of different temperatures, delivered by flow of thermal-controlled purified water, and chemesthetic stimuli. All thermal and chemical stimuli were tested on individual trials. In between trials, the mouth was continuously adapted to oral delivery of 35°C water, which intended to keep oral tissue at an approximate closed-mouth temperature for mammals (Green, 1986). Each trial was composed of a pre-stimulus, stimulus, and post-stimulus period. During the pre-stimulus period, the 35°C water adaptation rinse continued to bathe the oral cavity. Switching fluid flow to the stimulus solution started the stimulus period. Following this period, flow then switched back to the 35°C adaptation rinse to begin the post-stimulus period and finish the trial. Once a trial completed, stimulus passages of the delivery system were rinsed with purified water to neutralize system temperature.

Vc neurons were first tested to respond to the temperature stimuli, with temperature-controlled volumes of purified water delivered to the oral cavity in randomized order for each cell. For temperature trials, the pre-stimulus, stimulus, and post-stimulus periods were each 5 sec long. Seven temperatures were tested on discrete trials. Cool temperatures (those <35°C) were 7°C, 13°C [near a threshold for noxious cold in humans (Simone and Kajander, 1997; Rainville et al., 1999)], 21°C (cool, near laboratory “room” temperature), and 28°C [near the cooling activation threshold for TRPM8 (McKemy, 2007; Tajino et al., 2011)]. Noxious hot temperatures included 46°C [above the threshold for heat-activation of TRPV1 (Caterina et al., 1997; Caterina et al., 2000)] and 56°C. Finally, neurons were also tested with 35°C water, which was neutral/isothermal with the adaptation rinse. Cells recorded from TRPM8 deficient and control mice were also tested with 40°C water to assess sensitivity to oral warming, as below.

Each stimulus temperature stated here represents mean oral temperature achieved on all trials that tested this temperature, measured during the last 4 sec of the stimulus period. For each mouse, rinse and stimulus temperatures were continuously measured at the point thermal fluids entered the oral cavity from the stimulus delivery system. Temperature data were acquired using an IT-1E thermocouple (time constant = 0.005 sec, Physitemp Instruments, Clifton, NJ) embedded in the tip of the oral delivery tubing and a digital thermometer (BAT-12, Physitemp Instruments). Temperature data were sampled at 1 kHz by the data acquisition system (Li and Lemon, 2019); this facilitated real-time measurement of the temperature of oral fluid flow alongside cellular spiking data (Figure 1C).

After completing the temperature trials, Vc neurons were tested with chemesthetic stimuli including 35°C 1.28 mM L-menthol (MENT), 35°C 1 mM allyl isothiocyanate (AITC; mustard oil), 35°C 0.1 mM AITC, and room temperature 1 mM capsaicin (CAP). MENT is an agonist of the cold receptor TRPM8 (McKemy et al., 2002). AITC engages the nocisensors TRP ankyrin 1 (TRPA1) and also TRP vanilloid 1 (TRPV1) at high concentrations (Jordt et al., 2004; Everaerts et al., 2011). CAP is an agonist of TRPV1 expressed by nociceptive neurons (Caterina et al., 1997).

MENT and AITC were dissolved in purified water and presented to the oral cavity using our stimulus delivery system. For these trials, the pre-stimulus period was 5 sec, the stimulus period was 20 sec, followed by a 95 sec post-stimulus period. The order of MENT and AITC trials randomly varied across cells.

CAP was dissolved in a vehicle of 1.5% ethanol and 1.5% Tween 80 in purified water (Ellingson et al., 2009). Unlike MENT and AITC, CAP was uniquely brushed onto the rostral tongue at room temperature, as described (Li and Lemon, 2019). CAP trials were structured to include a 5 sec pre-stimulus period, 20 sec stimulus period, and a longer 95 sec post-stimulus period due to lasting neural effects of CAP. On a separate control trial, the vehicle-only solution for CAP was also tested by brushing this solution onto the tongue, in the same way CAP was applied (Li and Lemon, 2019). CAP tests were always performed last due to the lingering effects of lingual capsaicin on Vc neurons (Carstens et al., 1998).

For some neurons, if they remained isolated and showed baseline-level firing approximately 20 min following completion of the above tests, they were re-tested with a different set of controlled water temperatures presented orally. These included 6°C, 13°C, 18°C, 26°C, 30°C, 33°C, 38°C, 46°C, and 54°C, tested using a lower oral adaptation rinse temperature of 30°C. This aimed to study if mild warming from a lower adaptation temperature (for example, from 30°C to 33°C or 38°C) could excite Vc cells; neurons responsive to mild warming are comparably rare in trigeminal pathways (Poulos and Lende, 1970b, a; Yarmolinsky et al., 2016; Leijon et al., 2019). The order of the different temperatures was randomized, as the above.

### Identification of stimulation sites

At the conclusion of recording sessions, mice were euthanized by an overdose of pentobarbital sodium (≥ 130 mg/kg, i.p.) combined with 2% isoflurane. Mice were then transcardially perfused with 0.9% NaCl followed by a solution of 4% paraformaldehyde and 3% sucrose dissolved in 0.1 M phosphate buffer. Brains were removed and stored in a 0.1 M phosphate buffer containing 4% paraformaldehyde and 20% sucrose at ∼4°C. A sliding microtome (SM2010R, Leica, Deer Park, IL) was used to cut coronal sections (40 µm) of each brain, including forebrain tissue around the thalamic area to verify stimulating electrode sites. Sections were mounted onto gelatin-coated slides. The slides were Nissl stained, with electrode tracts compared to anatomical landmarks identified using a mouse brain atlas (Franklin and Paxinos, 2008).

### Data analysis and statistics

#### Neural sample

A total of 77 Vc units were recorded from wild-type mice (35 cells from female mice and 42 cells from male mice) and 39 Vc units were recorded from TRPM8-KO mice (21 from female mice and 18 from male mice). Sex failed to influence Vc cellular responses to thermal stimulation, with no significant sex × stimulus interactions (F_(2,159)_ = 0.082, *P* = 0.930, for wild-type mice; F_(1.6,58)_ = 0.585, *P* = 0.521, for TRPM8-KO mice; two-way ANOVA with *P*-levels corrected by the Greenhouse-Geisser method for lack of sphericity) or main effects of sex (F_(1,75)_ = 0.010, *P* = 0.919, for wild-type mice; F_(1,37)_ = 0.207, *P* = 0.652, for TRPM8-KOs). Thus, data from males and females were grouped and analyzed together. While in some cases multiple neurons were recorded from the same animal, the coordinates for these neurons differed from each other. Such cells were treated as independent units.

#### Measurements of neural activity

We calculated typical mean firing rate responses to temperature and chemesthetic stimuli and also temporal AP discharge patterns to thermal stimuli for Vc cells. Temporal measures were included because we observed that Vc neurons could display diverse transient (phasic) and sustained (tonic) AP response characteristics to oral temperatures *in vivo.* Diverse phasic and tonic responses also emerge in Vc neurons *in vitro* (Pradier et al., 2019), potentially reflecting physiological subtypes of Vc cells.

Temporal AP response patterns to temperatures were used to classify Vc neurons recorded form wild-type mice. To do this, hierarchical cluster analysis (HC; based on Ward’s method) was applied to Euclidean distance matrices computed among cells from their temporal spiking responses to 7°C, 13°C, 21°C, 28°C, 35°C, 46°C and 54°C. For each temperature trial, temporal responses were quantified as AP counts in 48 consecutive, non-overlapping 200 msec bins in the period from 5.4 sec (0.4 sec after stimulus onset to accommodate a brief initial artifact from the fluid control valve switching) to 15 sec (trial end). Temporal responses to temperatures for each unit were standardized prior to clustering by dividing the AP count in each bin by the standard deviation of the unit’s AP counts across all bins for the 7 temperature stimuli. Standardization aimed to facilitate clustering based on the shape of time-dependent AP firing rather than absolute firing amplitude to thermal stimuli.

Mean firing rates to oral temperatures were quantified as the average APs per sec that emerged during the stimulus period minus the mean APs per sec during the pre-stimulus period. As above, the response window was adjusted to 0.4 sec following stimulus onset.

Responses to MENT were calculated by taking the mean firing rate from stimulus onset to trial end minus the mean firing rate to isothermal water. This approach was used to accommodate the long and variable latency of Vc cellular responses to MENT.

Parametric and non-parametric analyses compared responses between different groups of neurons, as below. The organization of responses by Vc^Thal^ neurons to cooling temperatures (< 35°C) was studied using principal component (PC) analysis. Here, neural AP firing rates (in Hz) to oral thermal stimuli were standardized by division by the cell’s standard deviation of responses across the analyzed temperatures. PC analysis was applied using the *pca* function with default settings in MATLAB (ver. 2021a; MathWorks, Natick, MA) to linearly transform the original data set (standardized firing rates to oral temperatures × Vc^Thal^ cells) so that each PC provided a set of coordinates that captured variance in the data and was orthogonal to all other PCs. The structure and proximity of temperature responses plotted along PC coordinates reflected their organization.

#### Latency calculation and excitatory/inhibitory response

A Poisson-based algorithm (Chase and Young, 2007; Wilson and Lemon, 2014) was applied to AP trains to quantify the frequencies of Vc neurons in TRPM8-KO and wild-type mice that showed significant excitation or inhibition in firing to each temperature. In brief, to test the null hypothesis of no excitation, the algorithm estimated the probability that firing rates iteratively computed over sequential APs during the stimulus period were the same as the baseline firing rate in the pre-stimulus period. If this probability became lower than 10^−6^, the null was rejected and the time of the AP when this drop occurred was taken to mark the onset of a significant response and the latency of the neuron to respond to the temperature. If this criterion was not met for spikes falling within 4 sec of stimulus onset, the null was retained and the unit considered to not respond to the temperature. To test the null hypothesis of no inhibition, the procedure was the same except that the algorithm estimated the probability that firing rates iteratively computed over sequential APs the during pre-stimulus period were the same as the mean firing rate during the stimulus period; in this case, rejecting the null would reflect pre-stimulus firing that was greater than activity during the stimulus period. For both excitation and inhibition, the stimulus response window was considered to start at 0.4 sec following stimulus onset, as above.

#### Analysis of Vc neurons in TRPM8-KO and wild-type mice

To establish a baseline for analysis of the responses by the 39 Vc neurons recorded from TRPM8-KO mice, 49 of the 77 Vc cells sampled from C57BL/6J mice and tested with the same temperature series as the TRPM8-KO neurons were used as approximate wild-type control neurons. The Vc neurons recorded from TRPM8-KO mice were not all tested for thalamic projections; this factor was not assessed here. The analyzed neurons were tested to respond to oral delivery of water at 7°C, 13°C, 21°C, 28°C, 35°C, 40°C, 46°C, and 54°C following oral adaptation to 35°C water, as above; this series included 40°C to gauge potential responses to innocuous warming.

Features of thermal responses were compared between TRPM8-KO and wild-type Vc cells using inferential methods, as below. Further, relationships between the thermosensory tuning profiles of wild-type and TRPM8-KO Vc neurons were explored using multivariate and unsupervised machine learning techniques. PC analysis supported visualization of the distributions of Vc neurons from both mouse lines based on cellular AP responses (in Hz) to the thermal series. For this analysis, temperature responses were divided by the standard deviation of thermal firing by each neuron to standardize activity across cells.

To further compare neural distributions, TRPM8-KO and wild-type Vc cells were clustered using a technique based on non-negative matrix factorization (NNMF) of their AP responses (in Hz) to temperatures. NNMF is a matrix decomposition technique that can identify a small number of additive data features that compose patterns in a data set and can be used to classify data points (Lee and Seung, 1999; Brunet et al., 2004). We previously described use of this technique for classification of trigeminal-integrative neurons in the brain based on orosensory tuning (Li et al., 2022) and we followed that approach here. In brief, NNMF reduced a matrix of stimulus responses by individual cells to two non-negative, lower-dimensional matrices. These low-dimensional matrices partly weighted how well wild-type and TRPM8-KO Vc neurons fit into each of 2 or more clusters, denoted as *k*. Inclusion of cells in a cluster reflected a commonality in sensory responsiveness and tuning across oral temperatures. NNMF was performed using the *nnmf* function in MATLAB with default settings.

We found the optimal number of clusters by first repeatedly computing NNMF from random starting configurations at several sequential levels of *k*. We used *k* = 2 to 4 for classification of neurons via responses to five oral cooling/neutral temperatures (7°C, 13°C, 21°C, 28°C, and 35°C). The *nnmf* function was ran 100 times for each *k*. We discarded runs which converged to a rank lower than *k*, which was infrequent. The final *k* used to determine the number of cellular clusters present in the data was the largest *k* that resulted in the highest percentage of runs that repeatedly placed any given pair of neurons into the same or different cluster, evaluated using visual and quantitative methods (Brunet et al., 2004; Li et al., 2022). For the present analysis, the best *k* was found to be 2, indicating there were two major groups of TRPM8-KO/wild-type Vc cells based on thermal tuning.

Finally, we computed standardized Euclidean distances between responses to temperatures for wild-type and TRPM8-KO Vc neurons to assess similarities and differences in neural population responses to temperatures. Standardized Euclidean distances were computed using the *pdist* function in MATLAB. This standardization accommodated comparison of distances across mouse lines, which included different numbers of cells. Colormap matrices were used to visualize these distances.

#### General analyses

Confidence intervals for medians and means were bootstrapped using the MATLAB function *bootci* with 1000 resamples. Factorial ANOVA was used to analyze mean responses (in Hz) to temperatures. For ANOVAs involving repeated factors, Mauchly’s test evaluated the assumption that differences between group means were spherical. The Greenhouse-Geisser procedure was used to correct *F*-ratio degrees of freedom (df) if this assumption was violated. The corrected df were reported as real numbers, where applicable. Normality of data samples was addressed using the Jarque-Bera goodness-of-fit test in MATLAB. For group/factorial analyses, a natural log transformation was applied to all data points when data skew was present within groups. In some cases, this transformation normalized the data and parametric tests, such as ANOVA, were performed. When the transformation did not normalize skewed data, we chose to use non-parametric methods for data analysis. Log-transformed data were only used for analysis; the figures plot the original data points.

Statistical outcomes were evaluated using α = 0.05 corrected for multiple comparisons, when applicable, by using a false discovery rate control procedure (Benjamini and Hochberg, 1995). All analyses were performed in SPSS (version 27, IBM) and MATLAB using standard routines and custom code.

## RESULTS

### General response characteristics of cool-driven Vc neurons

Seventy-seven Vc neurons were recorded from C57BL/6J mice and tested using electrophysiological methods for antidromic responses from the VPM (thalamus). Fifty-two (68%) of these cells were successfully antidromically invaded from the VPM and classified as Vc^Thal^ neurons (e.g., Figure 1B), having axons that reached the thalamus. The remaining 25 neurons (32% of sample) failed the VPM antidromic response tests and were considered Vc^noThal^ cells that did not send axons to the thalamus. Across mice, the tip of the thalamic bipolar electrode predominantly targeted the ventromedial VPM (Figure 2).

**Figure 2.**
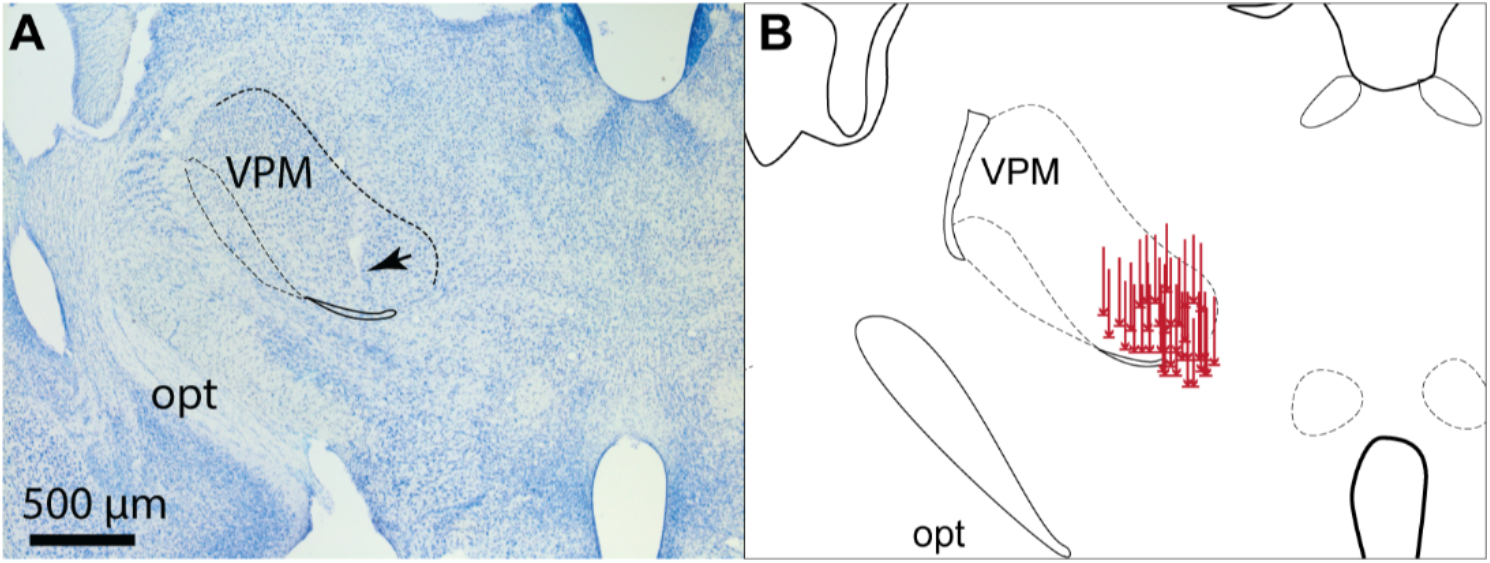
Reconstruction of VPM stimulating electrode sites. (A) Image of coronal brain section showing example electrode tract and tip placement (arrow) in the VPM in one mouse. (B) Drawing of stimulating electrode tract/tip locations in the VPM for 40 mice. Rostrocaudal dimension is conflated for simplification, opt, optic tract.

All Vc^Thal^ and Vc^noThal^ neurons analyzed here were sensitive to oral delivery of water at cool and cold temperatures below the 35°C oral adaptation temperature. Sixty-nine Vc neurons were tested for sensitivity to oral delivery of MENT, with most (*n* = 60, 87%) of these cells showing significant excitation to this cooling mimetic. No Vc cells showed significant excitation to oral stimulation with the nociceptive agents AITC or CAP. To address the potential for mild warming sensitivity, 13 of the 77 Vc units were also tested to respond to oral flow of water at 33°C and 38°C, and other temperatures, following whole-mouth adaption to 30°C. However, no cells showed a significant excitatory response to oral warming in this presumably non-noxious warmth range.

### Thalamic-projecting Vc neurons broadly respond to mild oral cooling and cold

Spontaneous activity did not differ between Vc^Thal^ and Vc^noThal^ neurons (non-significant main effect of group, F_(1,75)_ = 0.049, *P* = 0.826; Figure 3A), with a mean of 4.8 Hz for all cells. In contrast, responses to change in oral temperature were not the same across Vc^Thal^ and Vc^noThal^ neurons (significant main effect of group, F_(1,75)_ = 10.38, *P* = 0.002, observed power = 0.889). Specifically, Vc^Thal^ neurons included units that showed strong responsiveness to all cooling temperatures tested, from mild cooling to 28°C to the cold testing limit of 7°C. In contrast, Vc^noThal^ neurons were predominantly oriented to mild cooling temperatures, including 28°C and 21°C (Figure 3B). Post-hoc comparisons revealed that responses to 21°C (*P* = 0.0047), 13°C (*P* = 0.0003), and 7°C (*P* = 0.0002) were of greater magnitude in Vc^Thal^ neurons. Thus, certain differences were apparent in the electrophysiological response properties of the sampled Vc^Thal^ and Vc^noThal^ neurons, with broader responsiveness across cooling temperatures emerging in thalamic-projecting Vc cells.

**Figure 3.**
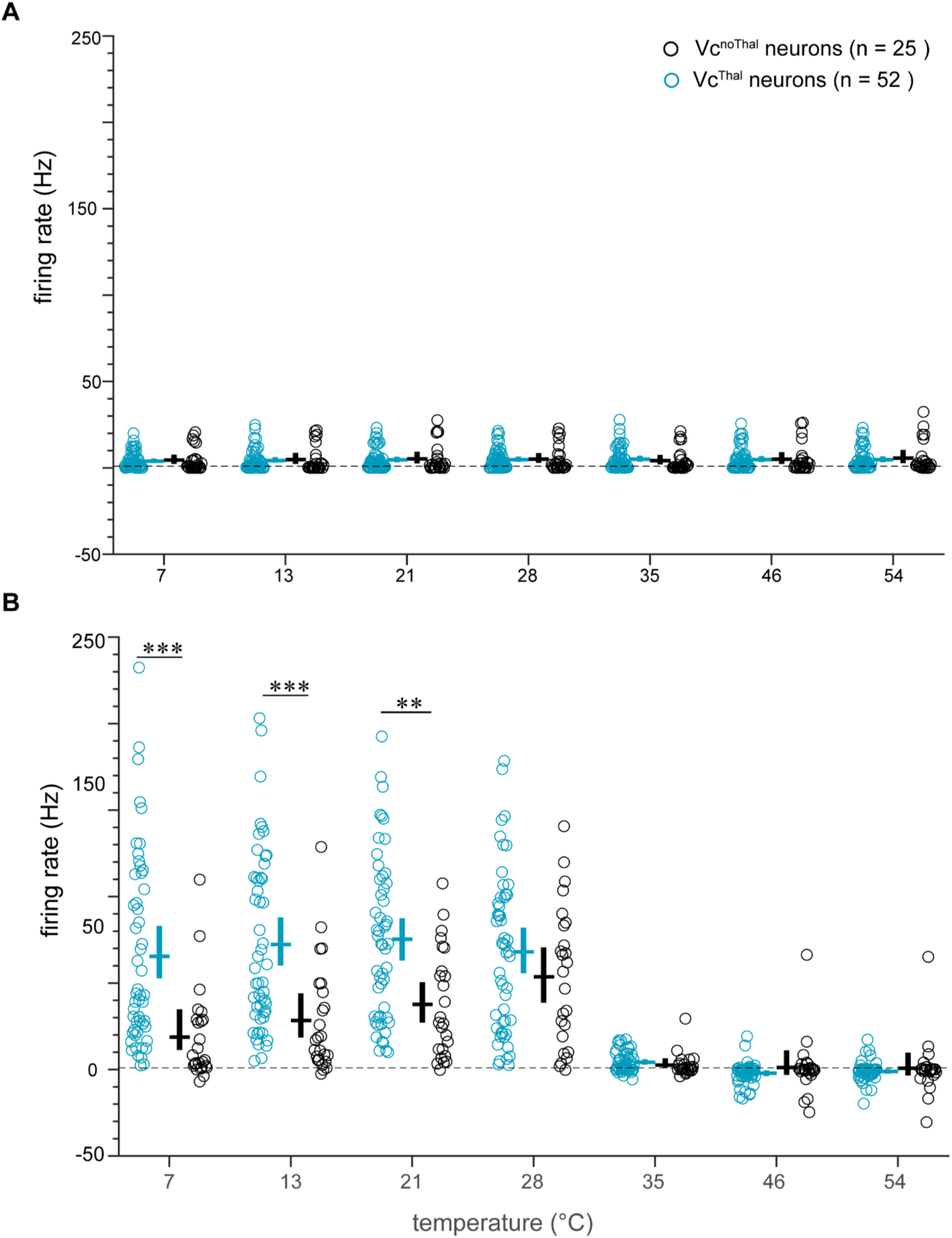
Vc neurons that project to the thalamus respond to a broad range of cool and cold oral temperatures. Shown are responses (In spikes per sec, Hz) during the (A) pre-stimulus and (B) stimulus periods of the temperature stimulation trials for Vc^Thal^ and Vc^noThal^ neurons (legend). Circles represent responses by individual neurons. Lines represent means (horizontal) and their 95% confidence intervals (vertical) for each distribution of responses. *\*\*, P <* 0.01 ; ***, *P <* 0.001.

### Diverse types of cool-driven Vc neurons project to the thalamus

Recent *in vitro* data show Vc neurons display heterogeneous temporal response patterns to depolarizing current injection. These patterns can appear as a phasic (rapidly rising then attenuating) AP discharge, tonic (sustained) firing, or also a delayed production of APs following depolarization (Pradier et al., 2019). This response diversity potentially reflects, in part, different physiological types of Vc cells. Here, we gauged sensory response diversity in Vc neurons *in vivo* by using HC to classify all Vc^Thal^ and Vc^noThal^ neurons by the time course of their AP discharge patterns to oral temperatures.

HC identified four major cell types across Vc neurons, with some types populating both Vc^Thal^ and Vc^noThal^ neurons and others predominantly arising in Vc^Thal^ cells. While neurons in all groups generally showed some degree of sustained firing during mild oral cooling to 28°C, cell types were distinguished by a characteristic decrease or persistence in AP discharge during later epochs of the thermal stimulus period as cooling temperatures approached and reached the cold limit of 7°C.

Two groups of neurons were classified as “phasic-cold” (groups 1 and 2; Figure 4A and 4B). Phasic-cold cells arose in similar numbers in Vc^Thal^ (group 1, *n* = 10; group 2, *n* = 7) and Vc^noThal^ (group 1, *n* = 11; group 2, *n* = 6) neurons and showed a transient increase in AP discharge to cold temperature drops to ≤13°C, followed by a rapid decrease in firing rate to near baseline levels for the remainder of the cooling period. These cells activated again, with a slow but pronounced rise and fall in AP discharge, once oral temperature returned to baseline (35°C) on cessation of cold stimulation (7° and 13°C); these were considered “off” responses, which were largely unique to phasic-cold cells. Considering response rates, Vc^Thal^ and Vc^noThal^ phasic-cold neurons

**Figure 4.**
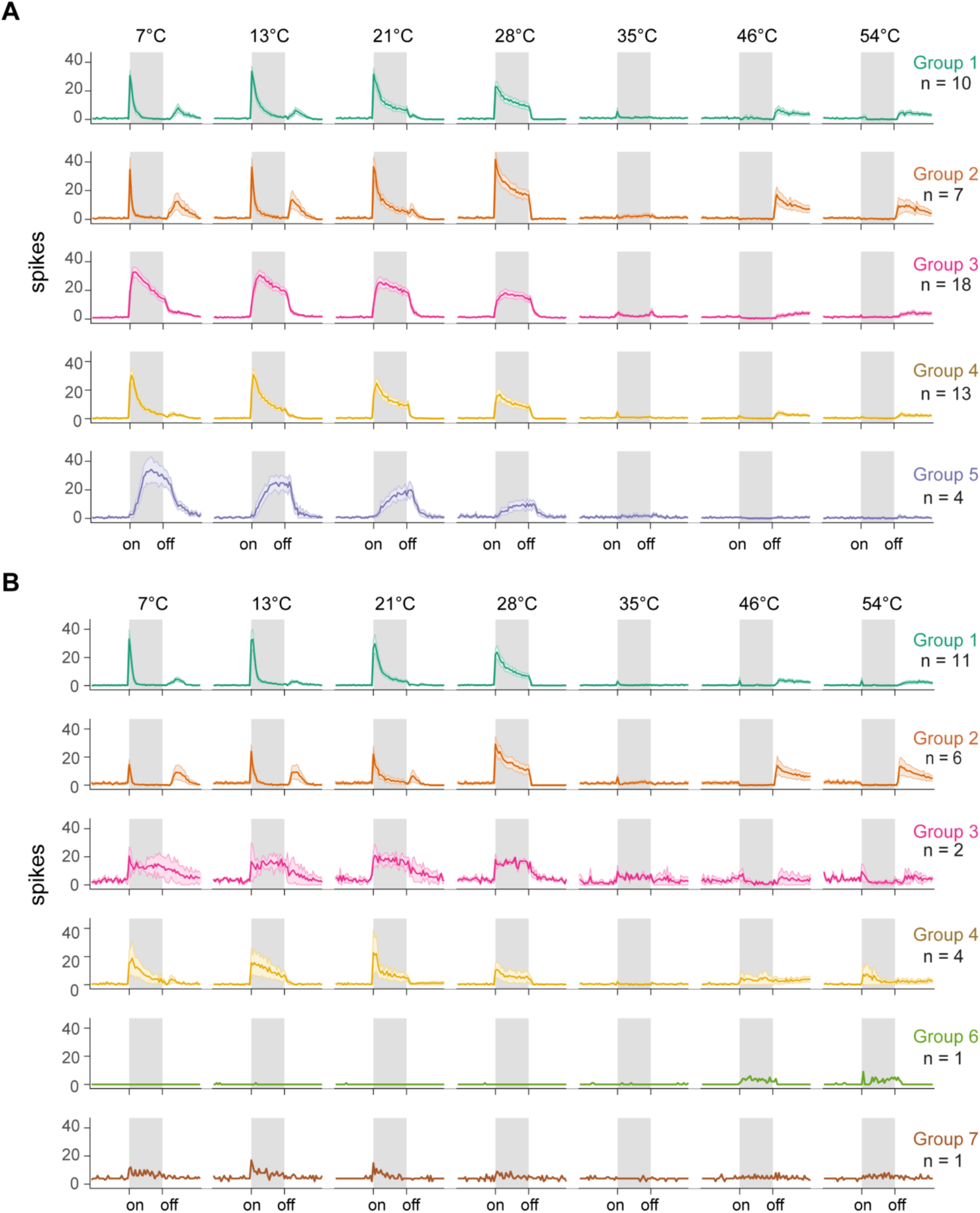
Diverse types of cool-driven Vc neurons project to the thalamus. Shown are temporal response kinetics to cool, neutral, and hot temperatures for HC-identified groups of (A) Vc^Thal^ and (B) Vc^noThal^ neurons. For each group, solid line tracks mean spikes per 200 msec during pre-, during- (grayed area between “on” and “off’), and post-stimulus periods. Shaded region surrounding mean line denotes standard deviation.

Neurons that showed comparably greater persistence to respond to cold temperatures were more common to Vc^Thal^ cells. These included group 3 neurons classified as “tonic-cold” (Vc^Thal^, *n* = 18; Vc^noThal^, *n* = 2), which generally showed a certain degree of continued firing for the duration of oral delivery of all cooling temperatures tested, including cold at 13°C and 7°C (Figure 4A and 4B). Accordingly, these neurons gave similarly strong response rates to cool and cold oral temperatures ≤ 28°C (Figure 5). Additionally, group 4 neurons (Vc^Thal^, *n* = 13; Vc^noThal^, *n* = 4) showed temporal responses to cold temperatures (13°C and 7°C) that appeared intermediate to phasic- and tonic-cold cells (Figure 4A and 4B). Group 4 neurons were broadly responsive to all cooling and cold temperatures tested (Figure 5). Finally, cluster analysis revealed one minor group of four Vc^Thal^ units that showed a slow, or “delayed”, increase in AP discharge to cooling onset (group 5 cells, Figure 4A). Delayed neurons, which did not emerge in Vc^noThal^ neurons, gave a monotonic rise in AP discharge over sequential cooling steps to reach a maximum response rate to cold at 7°C (Figure 5A).

**Figure 5.**
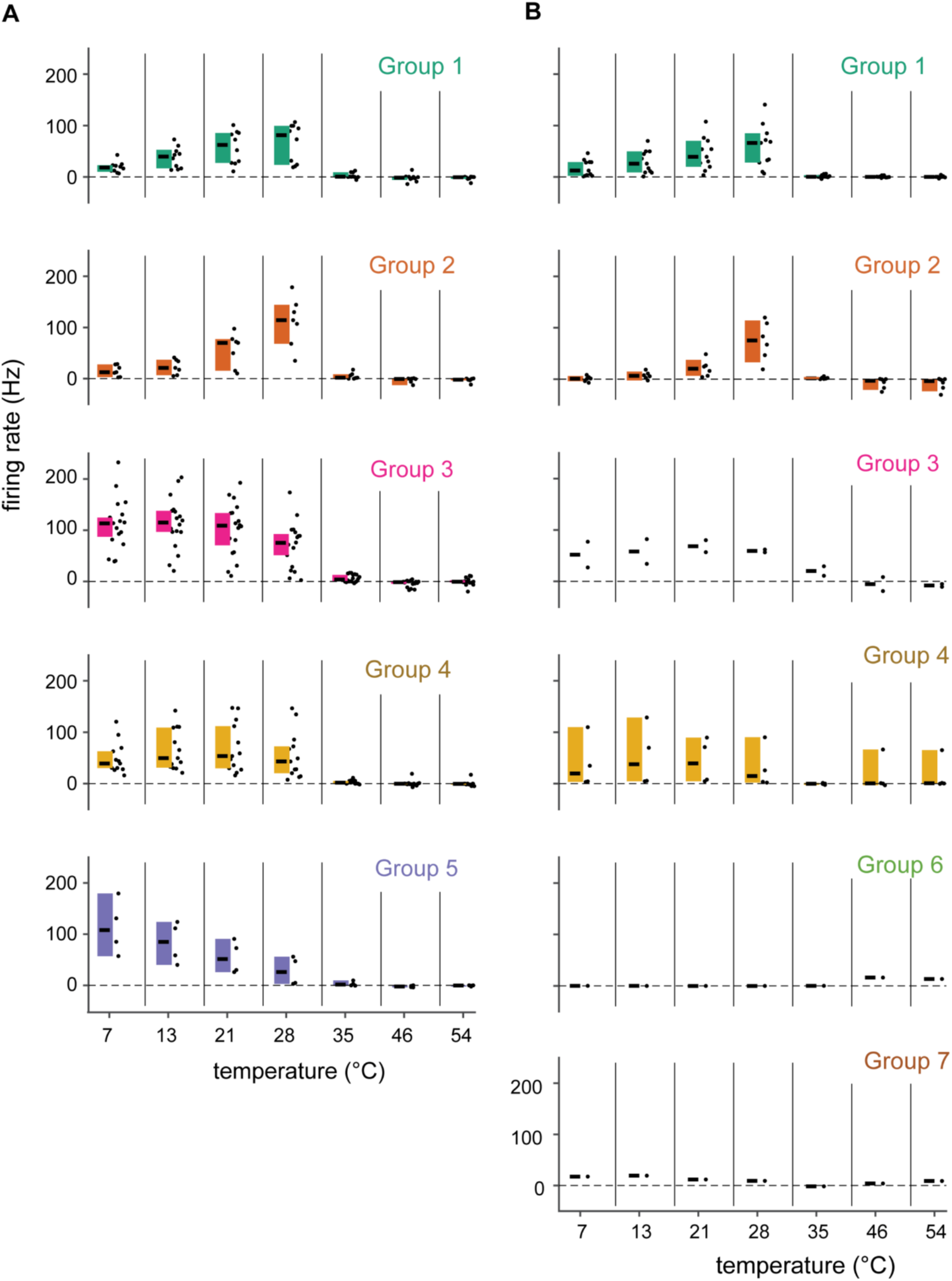
Cool-driven Vc neurons that project to the thalamus are tuned to different ranges of cooling temperatures. Points represent responses to each temperature (x-axis) by individual (A) Vc^Thal^ and (B) Vc^noThal^ neurons in the cell type clusters defined by HC (Figure 4). For each response distribution, horizontal line gives median and vertical bar represents the 95% confidence interval of the median.

These results reveal that cooling-driven Vc neurons that project to the thalamus compose a heterogeneous cellular population that includes neurons showing wide differences in response kinetics and tuning to mild oral cool and cold temperatures.

### Diversity among cool-driven trigeminothalamic neurons facilitates contrast between different oral temperatures

We applied PC analysis to firing rates to neutral and cooling temperatures by groups 1 through 4 Vc^Thal^ neurons to explore how these diverse types of thermosensitive cells may participate in oral temperature coding. We focused on only Vc^Thal^ neurons given their broader diversity and projections from the trigeminal medulla to the thalamus, potentially contributing thermosensory information to thalamocortical pathways. PC analysis reduced the high dimensionality of the original multivariate response data to lower-dimensional, visualizable representations that captured relationships between the patterns of activity evoked across neurons by temperatures. Separate PC analyses were applied to responses by individual cell groups and combinations of different groups. Scree plots of the variance explained by each PC assisted with understanding their relative contributions to the plotted data structures.

We observed that the first PC (PC1) accounted for a large majority the response variance to temperatures for Vc^Thal^ neurons in tonic-cold group 3, group 4, and phasic-cold groups 1 and 2 combined (Figure 6A, 6B, 6C; insets). However, there were some irregularities in the organization of responses to diverse temperatures by these cell types. Considering group 3 Vc^Thal^ neurons, responses to cool (21°C) and cold (7°C) were placed at similar coordinates along PC 1 and only marginally separated from mild cooling to 28°C (Figure 6A). This reflects the similarity of responses to all cooling temperatures displayed by group 3 cells (Figures 4 and 5) and implies these neurons, while responsive to temperature drops, may only poorly distinguish between cool and more intense cold temperatures. A similar clustering of responses to mild cool and cold was noted by PC analysis for group 4 Vc^Thal^ cells (Figure 6B). Finally, the responses of group 1 and 2 Vc^Thal^ neurons separated mild cooling to 28°C from other temperatures along PC1 but clustered the cold limit of 7°C together with 35°C (Figure 6C). This follows the similarly low responsiveness of these neurons to cold and warmth ≥35°C (Figure 5) and implies a graded response by these cells alone could not reliably track a thermal gradient.

**Figure 6.**
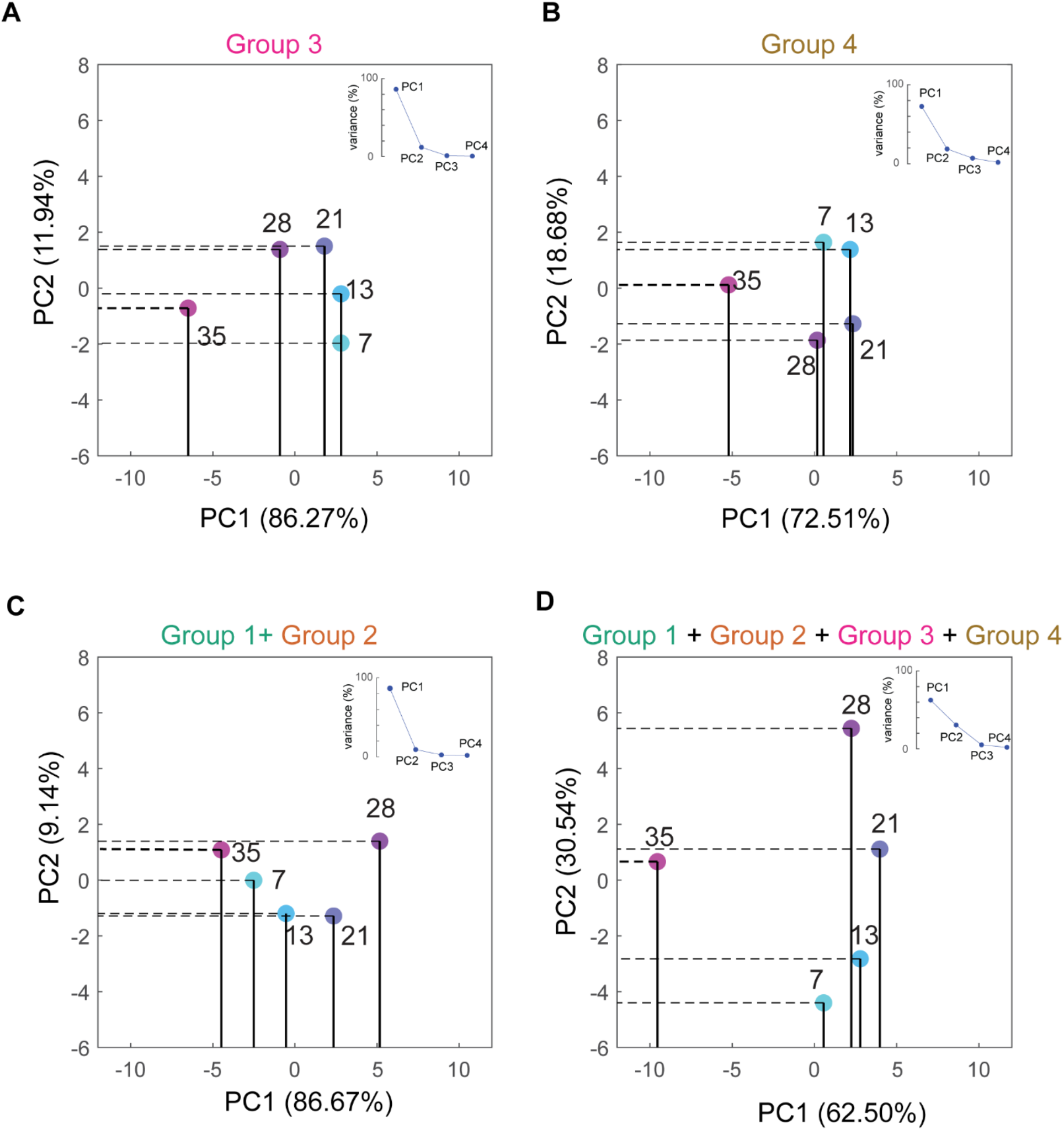
Diversity among cool-driven trigeminothalamic neurons facilitates contrast between different oral temperatures. Shown are PC analysis-recovered responses to neutral and cooling temperatures for individual (A, B), similar (C), and across diverse (D) types of Vc^Thal^ neurons. Cell types are from Figure 4A. Insets and parenthetical terms give variance explained by individual PCs. While individual (A, B) or like cell types (C) showed vagaries in their responses to, and arrangement of, the cool and cold temperatures, the combined response of diverse Vc^Thal^ neurons resulted in a systematic and ordered representation of the thermal series (D).

On the other hand, distinct cool, cold, and neutral/warm temperatures showed an orderly arrangement in two-dimensional PC space when the responses of Vc^Thal^ neurons in these four cell groups were combined together into a single population (Figure 6D). In this case, PC1 and PC2 both substantially accounted for thermal response variance (Figure 6D, inset). PC1 separated the population responses to oral cool and cold from neutral/warm 35°C. The degree of separation of warm from the cooling temperatures was larger than found for Vc^Thal^ cells in tonic-cold group 3, group 4, or in phasic-cold groups 1 and 2. In addition, responses to 28°C, 21°C, 13°C, and 7°C were systematically rank-ordered and spanned a uniquely broad coordinate range along PC2 (Figure 6D). These trends implied there was progressive and robust decorrelation between responses to diverse oral temperatures examined across the larger population of Vc^Thal^ neurons. In summary, the combined response of multiple diverse types of Vc^Thal^ cooling cells with different AP discharge kinetics and thermal tuning profiles offered greater contrast between representations for cooling temperatures than the actions of the cell groups alone.

### TRPM8 is required to establish Vc neurons tuned to mild oral cooling, but not intense cold

We recorded thermal responses from Vc neurons in mice gene deficient for the cooling and menthol receptor TRPM8 (TRPM8-KO mice), and C57BL/6J approximate wild-type control mice, to deduce the contribution of TRPM8 to oral temperature processing in central V circuits. Firing rates during the pre-stimulus period did not differ between TRPM8-KO (39 cells) and wild-type (49 cells) Vc neurons (Figure 9A, F_(1,86)_ = 1.268, *P* = 0.263). Almost all Vc neurons sampled from wild-type mice showed significant excitation to all cool (28° and 21°C) and cold (13° and 7°C) temperatures (Figure 7). Notably, we found that among TRPM8-KO Vc neurons, all or a marked majority (∼70% or more) showed significant excitation across all cool (28° and 21°C) and cold (13° and 7°C) temperatures (Figure 7), revealing Vc neurons maintain sensitivity to temperature drops inside the mouth in the absence of TRPM8. However, whereas the frequencies of Vc cells sensitive to 21°C, 13°C, and 7°C were similar between TRPM8-KO and wild-type mice (Fisher’s Exact Test, *Ps* > 0.05; Figure 7), significantly more wild-type cells were excited by mild cooling to 28°C (Fisher’s Exact Test, *P* = 0.001; Figure 7). This implies the loss of TRPM8 dampened V neural sensitivity to mild cooling, with less influence on colder temperatures.

**Figure 7.**
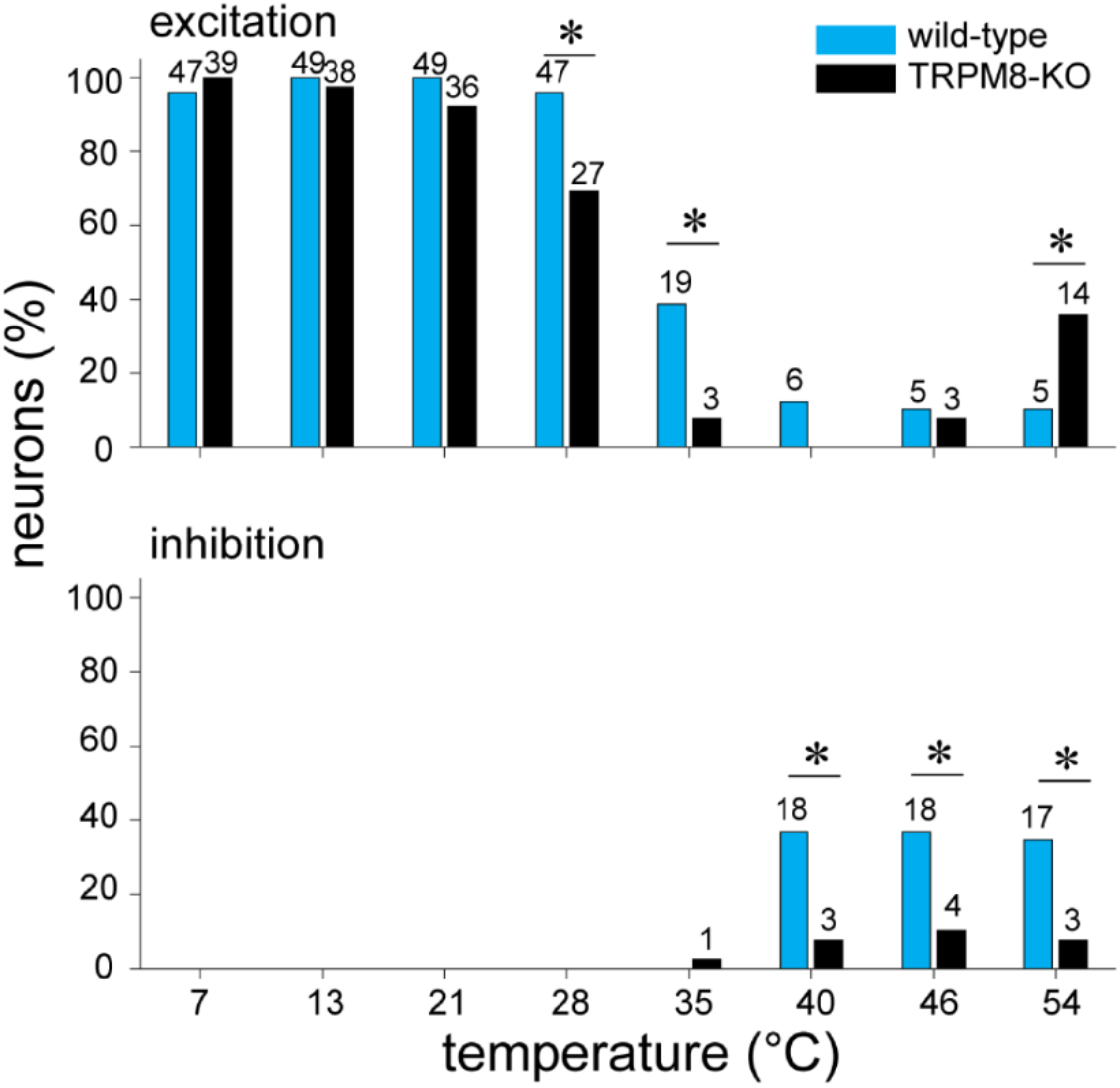
Vc neurons in TRPM8-KO and wild-type mice show differences in excitation (top) and inhibition (bottom) to oral temperatures. In both panels, bars give the percentage of neurons in each mouse line (legend) that showed a significant change in firing from baseline; the actual number of neurons is plotted above each bar. Data are from 49 wild-type and 39 TRPM8-K0 Vc cells. Significantly fewer TRPM8-K0 Vc neurons were excited by mild cooling (28°C; top panel). Moreover, fewer TRPM8-KO cells were inhibited by non-noxious warming (40°C) and noxious heat (46°C and 54°C; bottom panel). *\*, P <* 0.01.

**Figure 8.**
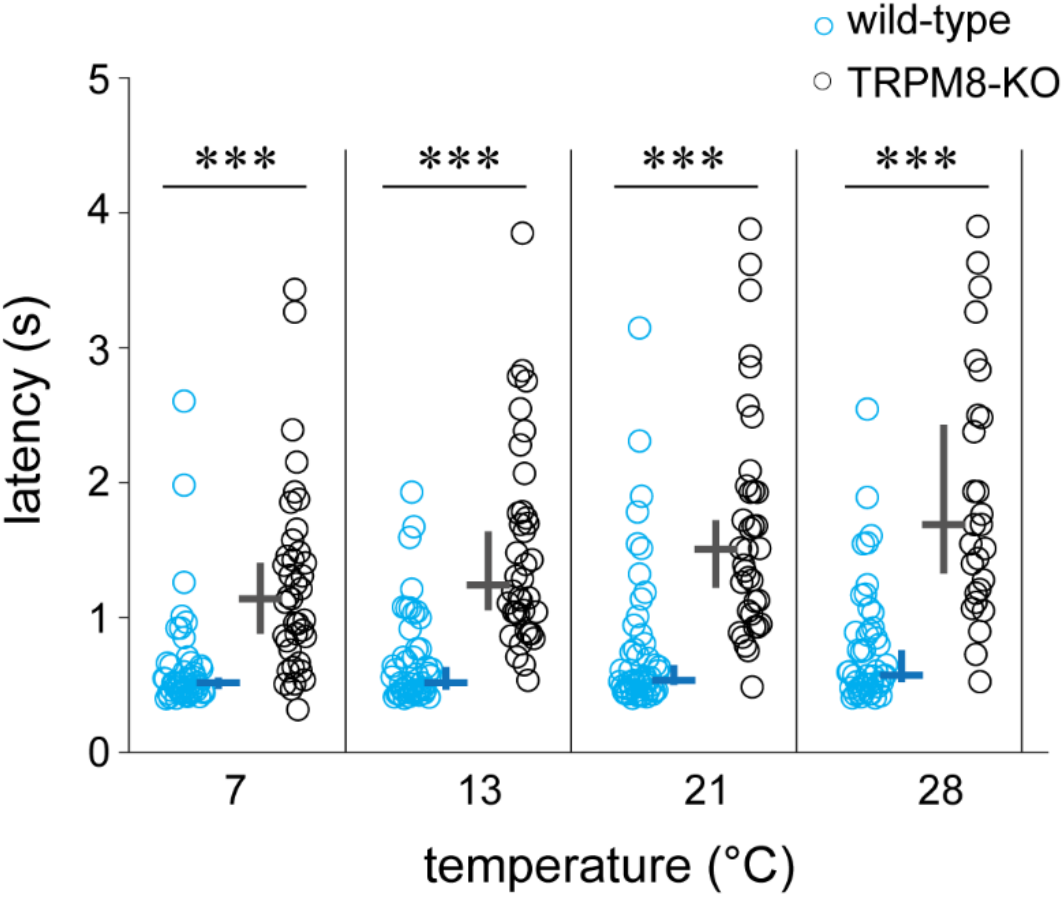
Vc neurons in TRPM8-KO mice showed slower-onset responses to cooling temperature stimulation of the oral cavity. Plotted are latencies to the first spike of the response to oral cooling stimulation for individual Vc neurons (circles) in TRPM8-KO and wild-type mice (legend). Lines represent medians (horizontal) and their 95% confidence intervals (vertical) for each latency distribution. ***, P< 0.001.

**Figure 9.**
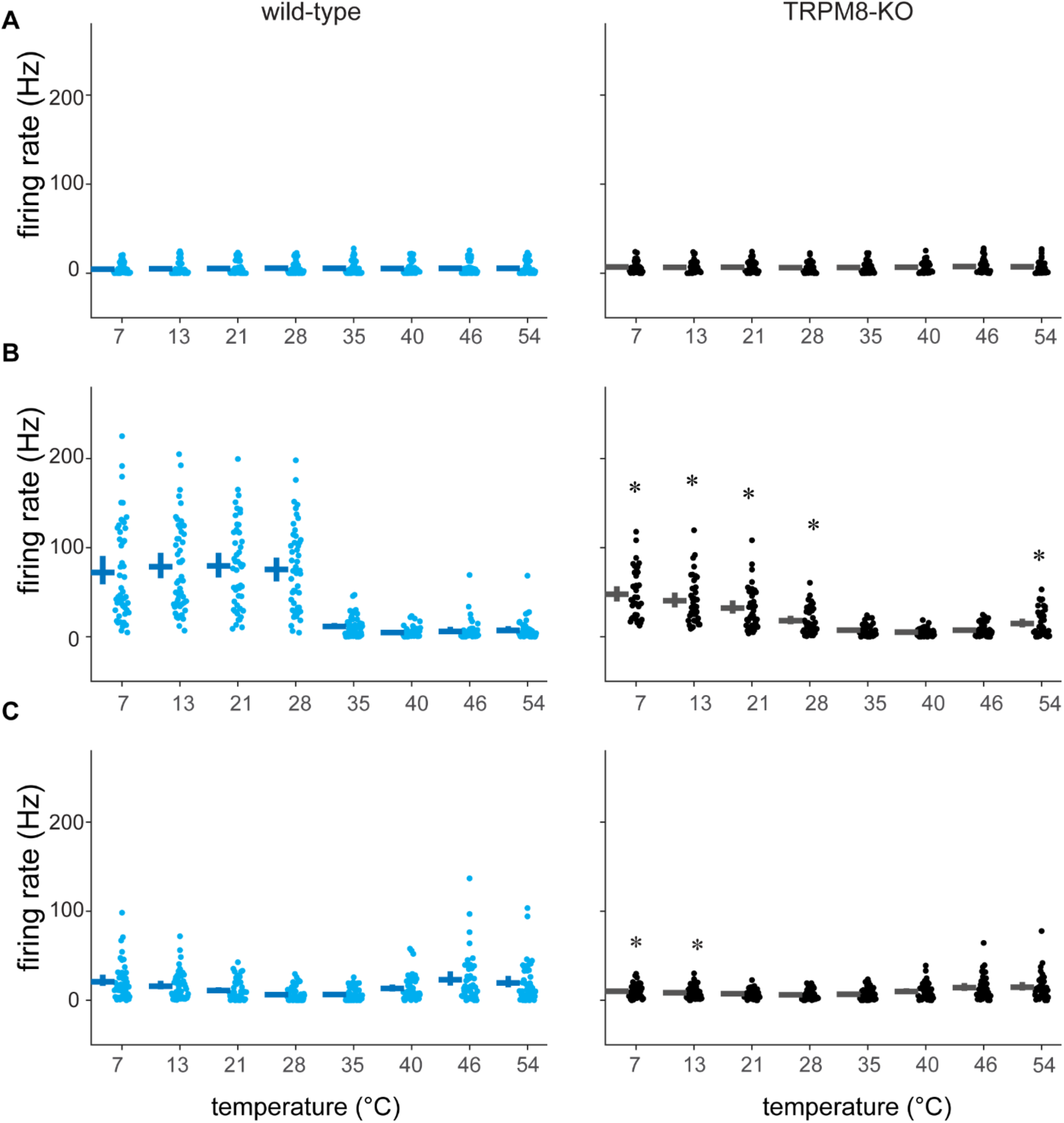
Vc neurons in TRPM8-KO mice showed reduced responses to cooling temperatures compared to control mice but maintain responsiveness to cold. Shown are firing rates (points) by individual Vc neurons in wild-type (left column) and TRPM8-KO (right column) mice during the (A) pre-stimulus, (B) stimulus, and (C) post-stimulus periods of temperature trials. Lines represent means (horizontal) and their 95% confidence intervals (vertical). *\*, P<* 0.01 for comparisons of temperature responses between wild-type and TRPM8-KO mice

Other differences in Vc thermal activity were also apparent between TRPM8-KO and wild-type mice. The median latency to significant response, when significant excitation was detected, was shorter in wild-type Vc neurons for all cooling temperatures (Wilcoxon rank sum tests with alpha corrected for multiple tests for 7°C, 13°C, 21°C, and 28°C, *Ps* < 0.001; Figure 8). This implied that in TRPM8-KO mice, the onset of Vc cooling responses arose at a slower pace. Further, the faster-onset responses to cooling temperature in wild-type mice happened at different latencies across all cooling temperatures but with small differences between medians (median values: 516 msec at 7°C, 515 msec at 13°C, 535 msec at 21°C, 572 msec at 28°C; Friedman’s ANOVA test, *P* < 0.001, *n* = 47). In contrast, Vc neurons recorded from TRPM8-KO mice showed substantially shorter response latencies to cold at 7°C (median: 1139 msec) than mild oral cooling at 28°C (median: 1689 msec; sign test with alpha corrected for multiple tests, *Ps* < 0.01, *n* = 21).

On average, the magnitudes of Vc neuronal responses to each cooling temperature (28°C, 21°C, 13°C, and 7°C) were larger in wild-type compared to TRPM8-KO mice (*Ps* < 0.005, simple comparisons of mouse line at each temperature; temperature × mouse line interaction: F_(1.3,112.8)_ = 34.3, *P* < 0.001; Figure 9B). The after-stimulus discharge (i.e., an “off” response) to 13° and 7°C was also greater in wild-type neurons (*Ps* < 0.008, simple comparisons of mouse line at each temperature; Figure 9C). Moreover, wild-type Vc neurons showed equivalent average response rates across all cool (28°C, 76 Hz [mean] ± 7 [s.e.m.]; 21°C, 80 ± 7) and cold (13°C, 79 ± 7; 7 °C, 72 ± 8) temperatures (non-significant simple effect of temperature, *P* = 0.23; Figure 9B). Yet in TRPM8-KO Vc cells, mean response rates to cool (28°C, 18 ± 2; 21°C, 32 ± 4) and cold (13°C, 40 ± 4; 7°C, 48 ± 5) increased with sequential cooling steps (significant simple effect of temperature, *P* < 0.001), with 7 °C eliciting the largest cooling/cold response in the absence of TRPM8 (Figure 9B). Altogether, the above data suggest that silencing TRPM8 produces a greater deficit on the responsiveness of Vc neurons to mild oral cooling than intense cold.

We used multivariate and machine learning techniques to compare the distributions of Vc thermosensory neurons between mouse lines based on thermal tuning. PC analysis was applied to sort all wild-type and TRPM8-KO Vc cells by their standardized firing rates to all oral temperatures. The first principal component (PC1), which accounted for a majority of the response variance (Figure 10A, inset), separated a subpopulation of wild-type neurons from a combined pack of wild-type and TRPM8-KO Vc cells (Figure 10A). Moreover, NNMF clustering of neurons by thermal responses recovered two clusters of cells, with one group a unique subpopulation of Vc neurons that largely overlapped with the wild-type units separated by PC analysis (NNMF-1; Figure 10B). This cluster was almost wholly populated by wild-type Vc cells (24 of 26 cells [92%]), with included neurons oriented to mild cooling at 28°C (Figure 10C). The second cluster of Vc neurons recovered by NNMF (NNMF-2; Figure 10B, 10C) included half of all wild-type Vc cells (25 of 49; 51%) and nearly all TRPM8-KO Vc neurons (37 of 39; 95%), with neurons in this cluster oriented to colder temperatures, including 13° and 7°C.

**Figure 10.**
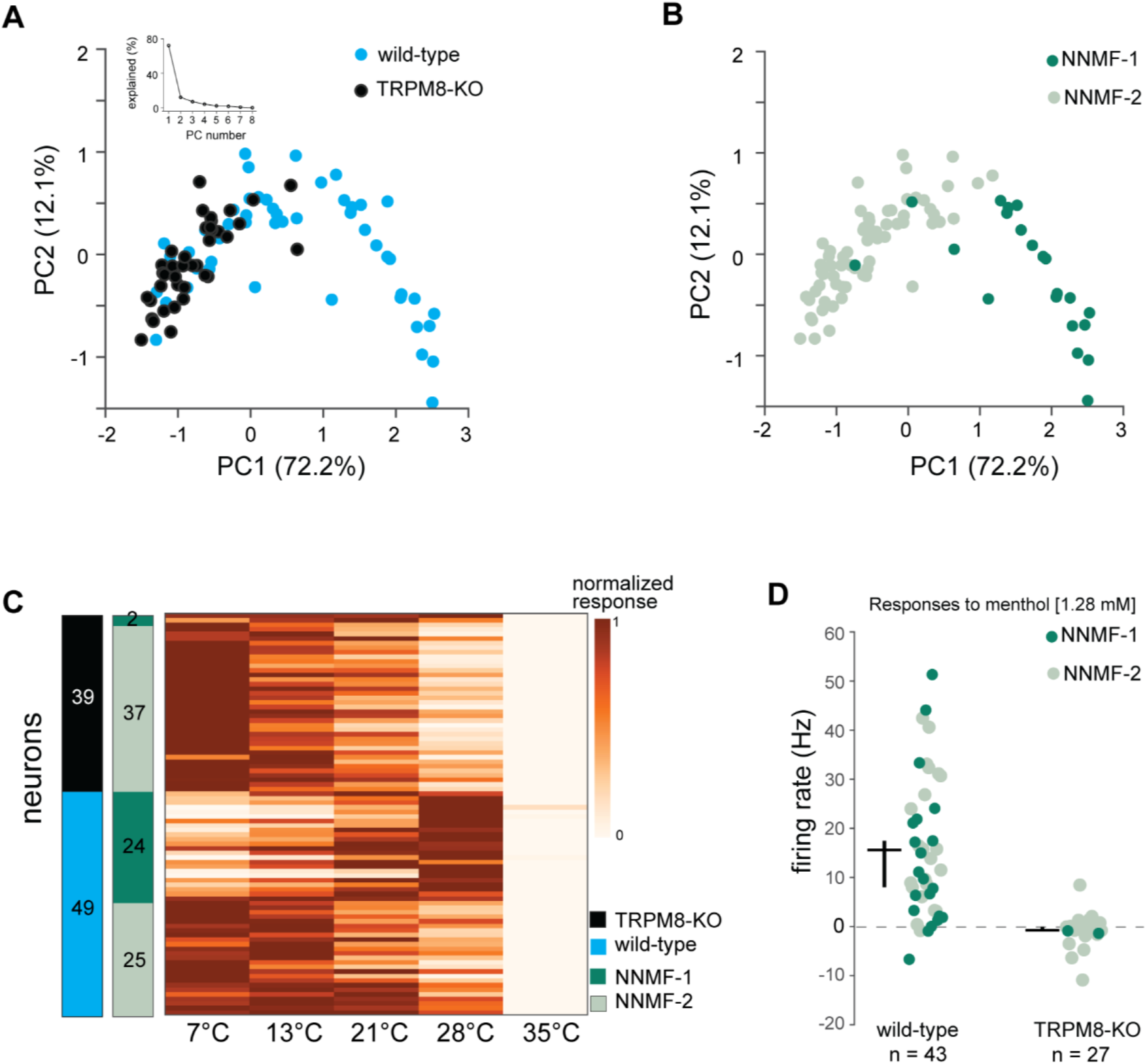
Vc neurons tuned to mild oral cooling are present in wild-type but missing in TRPM8-KO mice, which maintain Vc cells oriented towards intense cold. (A) Scatter plot of wild-type (n = 49) and TRPM8-KO (n = 39) Vc neurons (circles; see legend) recovered by PC analysis of thermal responses. Inset and parenthetical term gives variance explained by individual PCs. (B) Same as panel A except legend codes neurons based on their group identified by NNMF-based classification of cells by thermal responses. Note the unique group of wild-type neurons identified by both PC (cells ∼1 or more along PC 1) and NNMF (NNMF-1) analyses. These neurons were tuned to mild oral cooling (28°C) and unique to wild-type mice, as in panel C. (C) Normalized responses to neutral and cooling temperatures by TRPM8-KO an wild-type Vc neurons, divided into the two NNMF clusters. For each neuron, responses are normalized across temperatures to make the smallest response 0 and the largest response 1, as represented by the color map legend. (D) Responses to oral delivery of menthol in individual TRPM8-KO (n = 27) and wild-type (n = 43) Vc cells (circles). A loss of neural sensitivity to menthol was apparent in the knockout line.

Overall, these data suggest TRPM8 afferent input is not needed to establish Vc neurons responsive to intense oral cold temperatures, such as 7°C, as these cells persist in the absence of TRPM8. However, TRPM8 *is* necessary to define Vc neurons tuned to mild cooling temperatures, such as 28°C, as such cells are present in wild-type but notably missing in mice that genetically lack TRPM8. The loss neurons tuned mild cooling when TRPM8 was silenced appeared to impact neural representations for such temperatures, with increased similarity emerging between Vc neural population responses to distinct mild cooling (e.g., 28°C) and warm temperatures (≥35°C) in TRPM8-KO compared to wild-type mice (Figure 11). Thus, TRPM8 input is needed to distinguish mild cooling from oral warming in Vc circuits.

**Figure 11.**
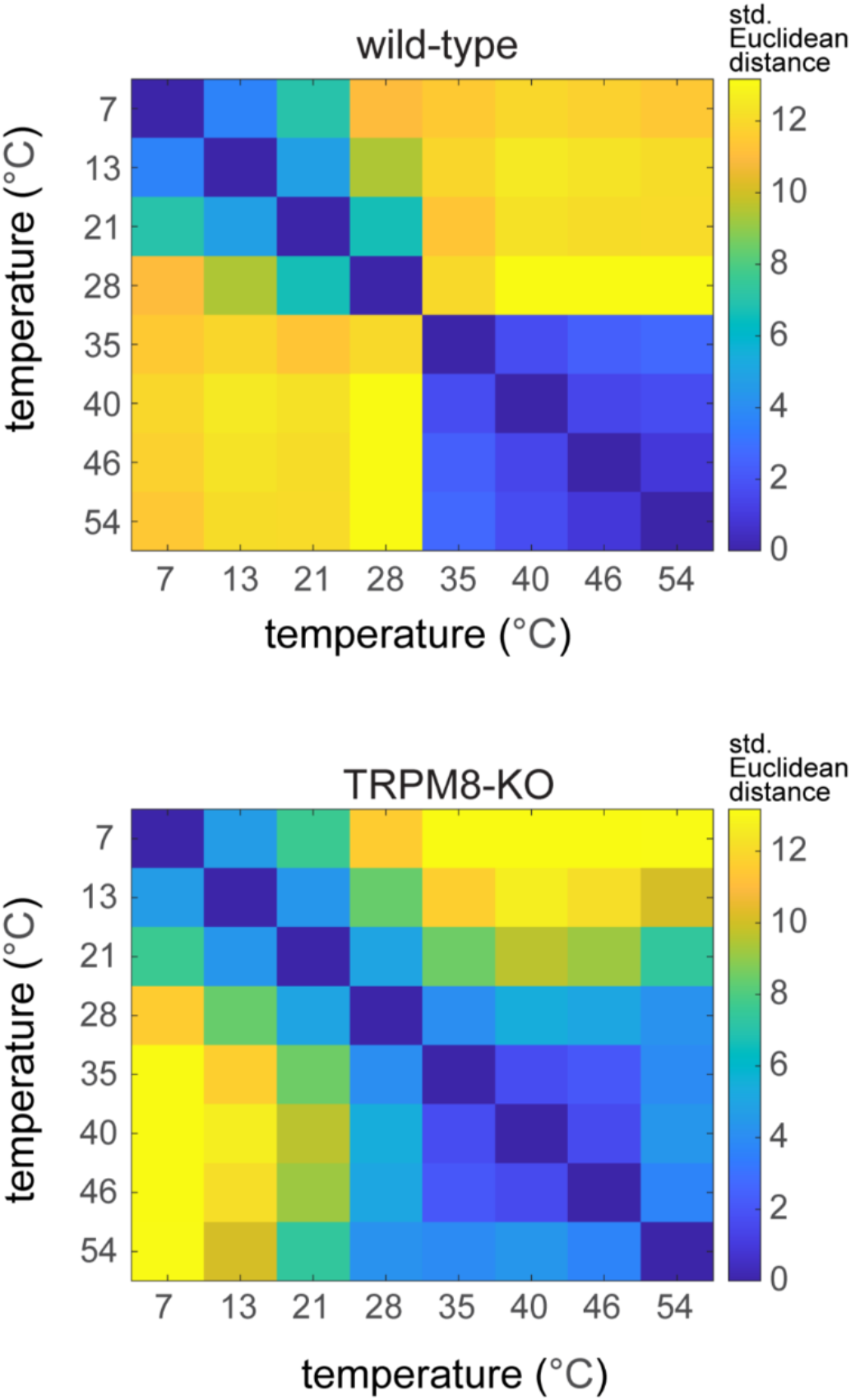
Silencing TRPM8 decreases neural contrast between cool and warm temperatures. Matrices plot standardized Euclidean distances (legend) between population responses to temperatures by Vc cells in wild-type (n = 49, top) and TRPM8-KO (n = 39, bottom) mice. Note the comparably increased similarity (i.e., lower distance) between Vc responses to mild cooling (28° C and 21°C) and warm (≥35°C) temperatures in TRPM8 deficient mice.

Finally, cool/cold sensitive Vc cells in wild-type mice showed strong responses to oral delivery of the cooling mimetic and TRPM8 agonist MENT, which failed to stimulate cold-sensitive Vc neurons in TRPM8-KO mice (Figure 10D), agreeing with their genotype.

### Silencing TRPM8 impacts Vc neural sensitivity to oral warming

There were few Vc neurons in wild-type and TRPM8-KO mice that showed significant excitation to oral warming and heat. However, a larger number of Vc neurons in TRPM8-KO mice were significantly excited by noxious hot water at 54°C (Fisher’s Exact Test, *P* = 0.005; Figure 7). Further, 54°C caused larger mean response rates in Vc neurons in TRPM8-KO mice (15 Hz [mean] ± 2 [s.e.m.]) compared to wild-type controls (7 Hz [mean] ± 2 [s.e.m.], *P* = 0.007, Figure 9B). More Vc units in wild-type mice showed significant inhibition in firing to warming and heating to 40°C and 46°C (Fisher’s Exact Test, *Ps* ≤ 0.006; Figure 7) and also oral delivery of intense noxious heat at 54°C (Fisher’s Exact Test, *P* = 0.004; Figure 7). Thus, the absence of TRPM8 associates with a change in sensitivity and responsiveness to oral heat in Vc cells.

Finally, significant excitation to both oral cold (7° or 13°C) and heat (46° or 54°C) was observed in 6 Vc neurons in wild-type mice and 14 Vc neurons in TRPM8-KO mice. The larger percentage of cold/heat “bimodal” thermosensory neurons in TRPM8-KO mice was significant (Fisher’s Exact Test, *P* = 0.005) and was also apparent when plotting mean responses to all temperatures for wild-type and TRPM8-KO Vc cells (Figure 12).

**Figure 12.**
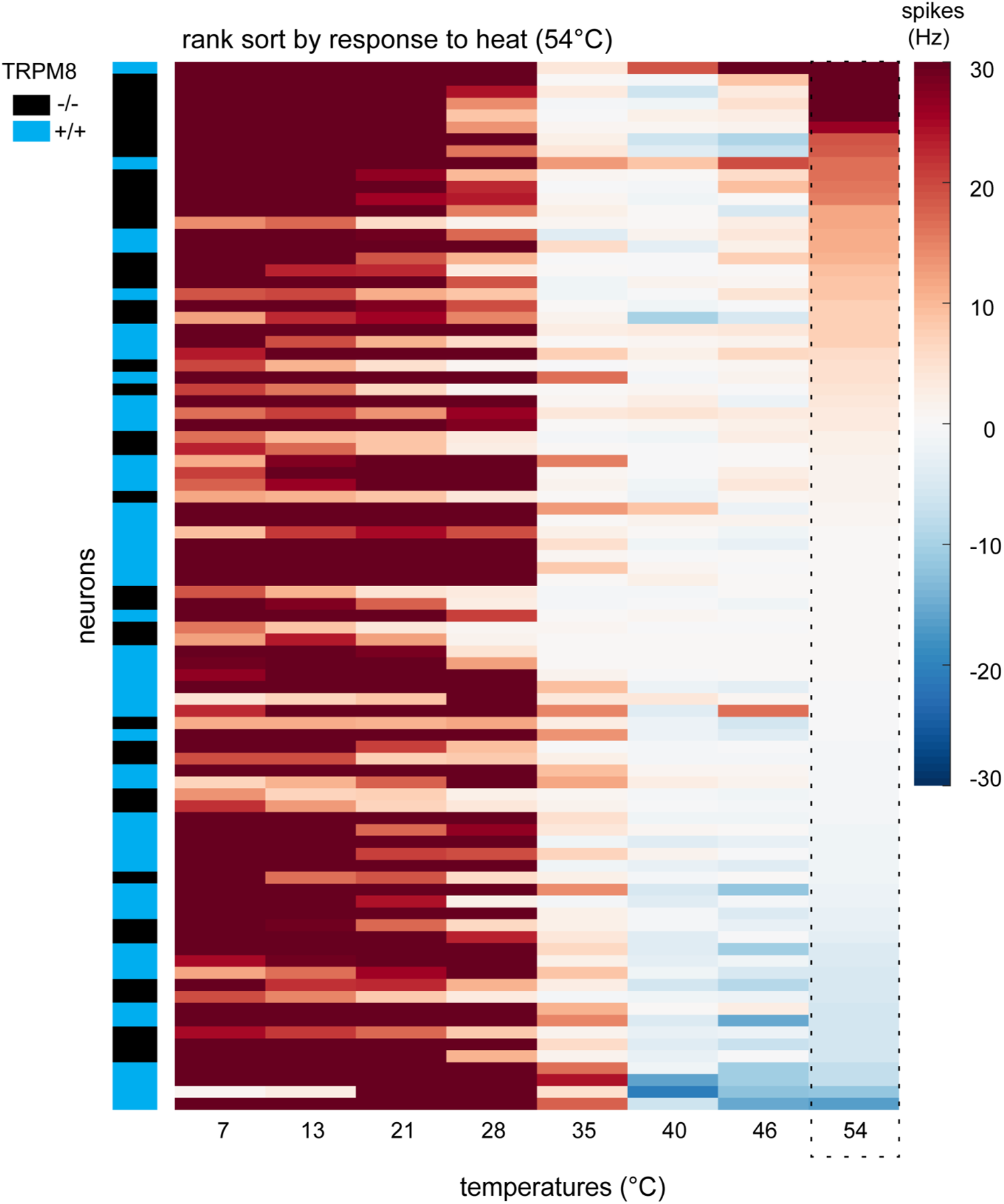
Bimodal (cold and hot excited) Vc neurons were more frequent in TRPM8-KO mice. Heatmap shows responses (in Hz, legend) to different temperatures for wild-type (+∕+) and TRPM8-KO (-∕-) Vc neurons (n = 88) rank-ordered by their response to 54°C.

## DISCUSSION

Our findings show that mouse Vc neurons that respond to oral cooling and project to the thalamus comprise heterogeneous cell types. These cell types were distinguished by differences in temporal AP discharge kinetics, and associated mean responses and tuning, across a broad range of mild cool (e.g., 28°C) to cold (e.g., 7°C) temperatures. Analyses of thermosensory information revealed that combining the responses of multiple, differently tuned types of these trigeminothalamic tract neurons offered greater contrast between representations of cool, cold, *and warm* temperatures compared to the responses of the individual neural types alone. Moreover, we found that Vc neurons oriented to mild cool oral temperatures critically depend on input from TRPM8 afferents. Vc cells tuned to mild cooling were frequently sampled in wild-type mice but were nearly absent in mice gene deficient for TRPM8. Notably, this absence of cooling neurons in TRPM8-KO mice decreased distinctions between Vc population responses to cool and warm temperatures ≥35°C. This implies TRPM8 input is needed to distinguish oral cooling from warming signals, agreeing with a role for TRPM8 in warmth recognition in other systems (Pogorzala et al., 2013; Paricio-Montesinos et al., 2020). While showing a deficit in sensitivity to mild cooling, TRPM8 deficient mice maintained Vc neurons responsive to and oriented towards cold temperatures such as 7°C, implying non-TRPM8 mechanisms contribute to Vc activity to cold stimulation of intraoral skin.

### Diverse types of Vc neurons signal oral cooling to the thalamus

Our recordings uncovered several types of Vc neurons in mice that respond to cool and cold stimulation of the oral cavity. These cell types were statistically identified based on differences in the time course of AP discharge to oral cooling temepratures. The majority of Vc^Thal^ neurons, which contributed to the trigeminothalamic circuit, comprised cell types that were generally considered phasic- or tonic-cold cells. Phasic-cold neurons showed only a brief transient (i.e., phasic) increase in AP discharge to oral cold (13°C and 7°C), with more sustained firing to mild oral cooling (21°C and 28°C) during the stimulus period (groups 1 and 2, Figures 4A). This temporal response characteristic to temperature associated with the mean overall tuning of phasic cold neurons to mild cool temperatures, with low average responses to cold (Figure 5A). On the other hand, tonic-cold neurons showed greater sustain in firing to oral cold and mild oral cooling (group 3 cells; Figure 4A), which reflected their broad mean responses across the cooling steps tested (Figure 5A). Notably, tonic-cold neurons were frequently identified among Vc^Thal^ neurons but were rare among Vc^noThal^ cells, implying tonic-cold cells may mostly comprise thalamic-projecting Vc units. Moreover, Vc^Thal^ cells also uniquely included a subgroup of neurons that showed delayed onset in firing to all cooling temperatures (group 5 cells, Figure 4A), with a monotonic rise in mean AP discharge over sequential cooling steps to reach a maximum response to cold at 7°C (Figure 5A). These delayed-onset cells were oriented to more extreme cold temperatures that are sometimes discussed as noxious cold (Simone and Kajander, 1997; Rainville et al., 1999).

The present finding that cool-driven trigeminothalamic tract neurons comprise subgroups distinguished by differences in temporal response kinetics to oral cooling and cold follows prior electrophysiological and functional imaging studies of primary (peripheral) V neurons. These works identified that cooling-excited V ganglion cells also display differences in phasic or steady-state/tonic temporal responses to cool and cold temperature stimulation of the tongue and oral cavity (Poulos and Lende, 1970b; Yarmolinsky et al., 2016; Leijon et al., 2019). In some cases, phasic and tonic responses to cooling ramps in V ganglion cells were shown to associate with the tuning of these neurons to mild cooling or more intense oral cold, respectively (Yarmolinsky et al., 2016), similar to the present results. Taken together, the prior peripheral and present central data reveal that subpopulations of V cells showing unique temporal response characteristics and tuning to different ranges of cool and cold mediate the neural coding of a broad range of oral cooling temperatures from mouth to the thalamus. While the AP response kinetics of V ganglion cells likely provide bottom-up drive to shape temporal responses to oral temperatures in Vc circuits, recent *in vitro* functional data show Vc neurons, themselves, can display transient (phasic), sustained (tonic), or delayed AP discharge patterns to depolarization (Pradier et al., 2019). Thus, Vc neurons have intrinsic physiological properties associated with their temporal discharge patterns to temperatures observed presently *in vivo*. Both bottom-up and central processing may shape the temporal response kinetics and thermal tuning of Vc cells that reach the thalamus.

In the present work, cooling temperatures were applied as temperature ramps that rapidly and continuously changed from 35°C to lower temperatures. On trials that targeted oral delivery of extreme cold at 7°C, the thermal ramp “passed through” the mild cooling temperatures that, on average, best excited phasic-cold neurons. It is possible that the transient excitation these neurons displayed to oral cold partly reflects their activation to the segment of this ramp that transitions through mild cool temperatures to reach 7°C. Thus, neurons that show this feature may be selectively tuned to *only* mild cooling, with their phasic excitation to cold due to sensitivity to the transition phase of the cold ramp. Nonetheless, temperature change on oral epithelia could, arguably, always involves a continuous ramp-like change, as heat exchange would presumably still gradually arise in resting temperature tissue even if a discreet thermal step could be applied.

### Combinatorial coding of oral cooling

Considering Vc^Thal^ neurons, our analyses uncovered that more information about cooling temperatures was conveyed by the combined response of different cooling-excited subtypes of these neurons compared to the responses of the subtypes alone. For example, phasic-cold neurons in groups 1 and 2 showed strong mean responses to mild cooling to 28°C and 21°C, but comparably low average firing to cold at 7°C (Figure 5A). Cold evoked a response in these neurons that was of similar magnitude to warm temperatures ≥35°C (Figures 6C), implying a graded response could not effectively signal a cooling gradient. Further, tonic-cold neurons in group 3 showed broad responsiveness across all cooling steps (Figure 5A), which provided only limited information about differences between cool and cold temperatures (Figure 6A).

However, when the responses of these, and other, cell types were combined, systematic and clear distinctions between mild cooling and extreme cold, and separation of these responses from warming, became apparent (Figure 6D).

This observation agrees with the idea that in V pathways, distinctions and transition across a broad range of oral cooling and cold temperatures are mediated by an ensemble, or combinatorial, neural code (Lemon et al., 2016; Leijon et al., 2019; Lemon, 2021); other data on V oral-cooling neurons also agree with ensemble coding (Yarmolinsky et al., 2016). The present results extend this possibility to Vc neurons that maintain axonal projections to the thalamus and potentially drive thalamocortical circuits for thermosensation. Notably, evidence supporting a neural ensemble code for cooling also emerges in spinal thermosensory circuits supplying the limbs and paws, where cool-driven cells compose subgroups that activate to unique ranges of cool and cold temperatures rather than following a simple graded response to temperature drop (Ran et al., 2016; Wang et al., 2018).

While the present results reveal that information about an oral cooling gradient is apparent in an ensemble V code, further studies are needed to understand if combinatorial coding is sufficient to represent oral cool and cold sensations. Along this line, it is noteworthy that while phasic neural activation to cold may reflect transient sensitivity to “pass-through” temperatures on a cooling ramp, as above, this temporal AP discharge characteristic could also provide information concerning and differentiating cooling and cold temperatures. Moreover, the “off” responses that phasic-cold neurons generated on cessation of cold stimulation (Figure 4) could also contribute time-dependent information concerning temperature change.

### TRPM8 is necessary to establish Vc neurons tuned to mild oral cooling, but not intense cold

We compared Vc neurons between wild-type and TRPM8 gene deficient mice to examine how the cooling and menthol receptor TRPM8 on trigeminal fibers (McKemy et al., 2002; Bautista et al., 2007) contributed to thermal responses in Vc cells. A major finding pertained to the role of TRPM8 in driving the selectivity of Vc neurons to cooling temperatures. Whereas wild-type Vc neurons generally included cells oriented towards mild oral cooling (28°C) or cold (7°C), only cold-tuned Vc neurons were maintained in mice gene deficient for TRPM8 (Figure 10A-C). Moreover, oral delivery of menthol strongly excited many wild-type Vc cooling neurons but was a poor stimulus for TRPM8-KO Vc cells (Figure 10D). Overall, these data reveal that TRPM8 afferents are necessary to establish Vc neurons oriented towards mild cooling stimulation of the oral cavity. Vc neurons tuned to more intense oral cold appear to, at least in part, rely on input from TRPM8-independent, menthol-insensitive thermoreceptors on trigeminal fibers.

Prior *in vitro* functional data demonstrate that V ganglion cells from TRPM8 gene deficient mice display a reduction of neurons responsive to mild cooling (e.g., 22°C) but maintain a subpopulation of menthol-insensitive cells responsive to cold (e.g., 12°C) (Bautista et al., 2007). Additional *in vitro* data show that variable expression levels of, in part, TRPM8 associate with the thresholds for cooling temperatures to stimulate cultured V ganglion cells, with cellular excitation to small temperature drops and lower temperatures associated with elevated and reduced TRPM8 expression, respectively (Madrid et al., 2009). These results imply that TRPM8 is a strong driver of V afferent sensitivity to moderate cooling temperatures. This agrees with the present discovery that central Vc neurons tuned to mild oral cooling depend on TRPM8 input.

Mechanisms that contribute to Vc cellular responses to more intense oral cold in TRPM8 deficient mice remain unknown. It is tempting to speculate that these responses are contributed by excitation of TRPA1, which is expressed on V afferents (Jordt et al., 2004; Kobayashi et al., 2005) and has a role in orosensory avoidance of cooling mimetics (Lemon et al., 2019) in addition to cold sensing and nociception under certain conditions (e.g., Kwan et al., 2006; Kwan and Corey, 2009; Bernal et al., 2021; Yamaki et al., 2021). This would agree with a role for TRPA1 in mediating Vc neural sensitivity to cold temperatures such as 7°C. However, there are multiple complexities surrounding the study of thermal activation of TRPA1 (Kwan and Corey, 2009; Talavera et al., 2020) and more work is needed to directly address mechanisms of TRPM8-independent responses to oral cold in mouse Vc cells. Additional potential mechanisms were recently described (Bernal et al., 2021).

Genetic silencing of TRPM8 influenced several aspects of the thermal responses that remained in Vc neurons in TRPM8-KO mice, including increasing cellular latencies to respond to oral cooling and cold temperatures (Figure 8). Speculatively, this may reflect that TRPM8-indpendent signals concerning oral cooling traverse V afferents that conduct more slowly than those expressing TRPM8, or that TRPM8 deletion reduces summation of cooling afferent input onto Vc cells, delaying AP generation. Moreover, the loss of neurons tuned to mild cooling to 28°C (Figure 10A-C), and the substantially reduced responsiveness of Vc cells to 28°C and 21°C (Figure 9B), in TRPM8-KO mice dampened distinctions between neural population responses to moderately cool and *warm* temperatures ≥35°C (Figure 11). This suggests that TRPM8 input is needed to enable V neurons and networks to register differences between oral cooling *and oral warming*, and that TRPM8-KO mice may experience difficulties in behaviorally distinguishing these temperature conditions. Relatedly, mice gene deficient for TRPM8, or with acute inactivation of TRPM8 channels, display impaired behavioral performance for reporting differences between non-noxious warming (42°C) and lower temperatures (32°C) on the paw compared to TRPM8-intact mice, revealing TRPM8 signaling participates in warmth detection in spinal pathways (Paricio-Montesinos et al., 2020).

Our neurophysiological approach searched for but found only sparsely few Vc neurons excited by non-noxious oral warming in wild-type mice (Figure 7). This agrees with the rarity of cells excited by warming shown in other studies of V oral thermal processing (Poulos and Lende, 1970b, a; Yarmolinsky et al., 2016; Leijon et al., 2019). Notably, significantly more Vc neurons in wild-type mice showed inhibition, where AP discharge fell significantly below baseline firing rate, to non-noxious warming and noxious heating of oral tissue compared to TRPM8-KO Vc cells, which rarely displayed inhibition to warm temperatures (Figure 7). This suggests oral warmth inhibits Vc neurons through a TRPM8 mechanism, which may participate in signaling oral warming. Relatedly, mouse behavioral recognition of non-noxious warming on the paw is tied to warm-evoked inhibition of TRPM8-positive cool-driven cutaneous afferent fibers, which are missing in TRPM8-KO mice (Paricio-Montesinos et al., 2020). Further work is needed to address how behaviors guided by cool *and warm* temperature sensing inside the mouth are contributed by TRPM8-positive V afferents, and downstream Vc cells.

TRPM8 was knocked-out in the embryo in the TRPM8-KO mouse model used here. Thus, some degree of compensatory change may have contributed to the oral thermal responses we observed in Vc neurons in these animals. Along this line, we found that in TRPM8-KO mice, there were significantly more Vc cells excited by brief oral delivery of noxious heat at 54°C (Figure 7), and more neurons displaying bimodal responsiveness to intense oral cold and heat (Figure 12), compared to wild-type mice. This may reflect that TRPM8 afferent signals shape the development and balance of sensitivity to oral cooling and heating displayed by V neurons and circuits. Further delineation of how TRPM8 signaling impacts the functional organization of V pathways, and defining how TRPM8-dependent and -independent neural messages influence orosensory-guided behaviors towards temperatures, will shed deeper light on how the brain processes and represents oral thermosensory information, and how oral thermal coding compares to spinal thermosensation.

## Acknowledgements

Supported by NIH grant DC 011579 to C.H.L.

